# Graph deep learning reveals multiple signal pathways activated in anti-citrullinated protein antibodies stimulated synoviocytes

**DOI:** 10.1101/2022.03.15.484500

**Authors:** Meng Sun, Yang Chen, Petter Brodin, Anca Catrina

## Abstract

We present a study of anti-citrullinated protein antibodies stimulated fibroblast-like synoviocytes from patients with rheumatoid arthritis, which is powered by a novel graph deep learning framework applied to single-cell mRNA expression data. This new analytical framework discovered dominant pathways that suggest IL1-IL1R mediated signaling, a novel signal transduction response to ACPA-stimulation, and comfirmed our previous finding of PI3K/AKT activation in response of ACPA-stimulation as well. The study demonstrated the capability of graph deep learning framework on signaling pathway analysis. The findings suggest that ACPAs contribute to distinct pathogenic events through activating multiple signaling pathways.

## 1 Introduction

Anti-citrullinated protein antibodies (ACPAs) are presence in a majority of patients with rheumatoid arthritis (RA) and are specific for this disease. Klareskog et al. [2009] ACPAs consist of a group of antibodies with different specificities towards citrullinated antigens and have been suggested to contribute to joint pain and bone loss already before onset of joint inflammation in RA. Bos et al. [2010], Kleyer et al. [2014], Krishnamurthy et al. [2019], Rantapää-Dahlqvist et al. [2003]

We and others have shown that ACPAs bind to the surface of developing osteoclasts (OC) and suggested that reactivity to citrullinated targets increase OC differentiation and bone loss. Krishnamurthy et al. [2016], Harre et al. [2012] Furthermore, in the synovial tissue, fibroblast-like synoviocytes (FLS) contribute to an inflammatory stroma that promote and amplify tissue-specific immune activation through the release of various cytokines and are able to grow into the cartilage surface, create an erosive interface by producing tissue remodeling proteases. Müller-Ladner et al. [1996], Lefèvre et al. [2009], Müller-Ladner et al. [1996] We have also shown that ACPAs binds to citrullinated targets on FLS and promote migration capacity via activating Akt phosphorylation at the Thr308 residue linked to PI3K activation. Sun et al. [2019] However, the potential mechanisms of ACPAs remains largely unknown, a complex scenario requests deep investigation in signaling transduction.

Challenge of signaling transduction investigation comes from complexity of the networks, which consists of sequences of proteins and molecules that transmit stimulation from outside of cell to inside target molecules. Existing methods rely on literature analysis and experimental confirmation, which is, however, highly time consuming and dependent on prior knowledge. In the present report, we design a framework to efficiently investigate multiple signaling pathways simultaneously based on Graph Neural Network (GNN) Gori et al. [2005], Scarselli et al. [2009]. GNN or Graph Deep Learning is a class of neural network that was introduced to deal with data best represented by graph data structures, for example biological networks. Inspired by this method, we design a graph data structure for single cell RNA sequence (scRNA-seq) profiled RA patients tissue derived FLS, and employ GNN based on the single cell data to discovery pathways activation between ACPAs and non-ACPA control IgGs (IgG) stimulated groups.

## 2 Material

### 2.1 FLS isolation

FLS were isolated from synovial tissues through enzymatic digestion using 4mg/ml Collagenase A and 0.1mg/ml DNase I (Roche, Mannheim, Germany). The dissociated cells were cultured in Dulbecco’s Modified Eagle Medium (DMEM) complemented with 10% heat-inactivated fetal bovine serum (FBS), 100U/mL penicillin, 0.1mg/mL streptomycin and 2mM L-glutamine (all from Sigma-Aldrich, St. Louis, MO, USA). Non-adherent cells were removed after overnight incubation and new medium was added.

### 2.2 Sample preparation

For 10x Genomics, FLS from 2 donors were prepared. Cells were cultured in T175 flask up to 80% confluence. Cell were starved by culture medium without FBS for 2 hours, followed by incubated with 2% FBS culture medium contains 1*μ*g/ml of either ACPAs or IgG for 6 hours. Single cell suspensions were collected with Trypsin-EDTA (Sigma-Aldrich) and passed through a 70*μ*m cell strainer.

For immunoblotting, FLS from 3 donors were prepared. Cells were culture in 6 well plates up to 80% confluence. Cells were starved by culture medium without FBS for 2 hours, followed by stimulation using 10*μ*g/ml of either ACPAs or IgG in 2% FBS culture medium up to 6 hours. Cells were collected by scraping at 2, 4 and 6 hours with leammli buffer, respectively.

### 2.3 RNA sequencing

Single-cell ATAC expression libraries for FLS were prepared by strictly following the protocols of the Chromium Single-Cell ATAC Library kit (10x Genomics). All resulting libraries were sequenced on the Illumina Xten PE-150 platform (ANOROAD and Novogene, Beijing, China). Raw 10x Genomics sequencing data were preprocessed with normalization, log transformation and standardization by using python libraries SCANPY Wolf et al. [2018] and Scikit-learn Pedregosa et al. [2011].

### 2.4 Immunoblotting

Proteins were collected from FLS using laemmli buffer with dithiothreitol. Cell lysates were separated by 4-20% sodium dodecyl sulfatepolyacrylamide gel electrophoresis (SDS-PAGE,Mini-PROTEAN TGX Precast Gels, bio-rad, Sweden) at 180V for 45 minutes. Protein were transferred onto nitrocellulose membranes (bio-rad, Sweden) at 25V for 30 minutes. The membranes were blocked with 5% non-fat milk (diluted with TBS containing 0.1% Tween-20) for 30 minutes at room temperature and incubated with primary antibodies overnight at 4°C.

The following antibodies were used: mouse anti-IL1R(1:1000, Abcam), rabbit anti-FAK (1:1000, Cell Signaling Technology), rabbit anti-Fizzled5 (1:1000, Cell Signaling Technology), rabbit anti-Wnt5a/b (1:1000, Cell Signaling Technology) and rabbit anti-GAPDH HRP (1:1000, Cell Signaling Technology). After washing, membranes were incubated with horseradish peroxidase (HRP)-conjugated goat anti-rabbit or anti-mouse IgG secondary antibody (1:5000, GE Healthy) for one hour at room temperature. Proteins were detected using SuperSignal West Femto Maximum Sensitivity Substrate (Thermo Fisher, USA). Images were captured by ChemiDoc MP Imaging System (bio-rad, Sweden)

## 3 Method

We propose the method for discovering signaling pathway activation by using the framework illustrated in Figure 1. It models a deep neural network to classify the sample data profiled by single cell sequencing against their stimulation groups. The model is trained by using a graph representation of the data structured based on known signaling pathways. Moreover, the model contains a layer applied with the attention mechanism (Veličković et al. [2018]) that learns the weights of how pathways contribute to classification. Afterwards, a higher weighted pathway indicates the higher chance of been activated, which can be used for further experimental confirmation.

**Figure 1:**
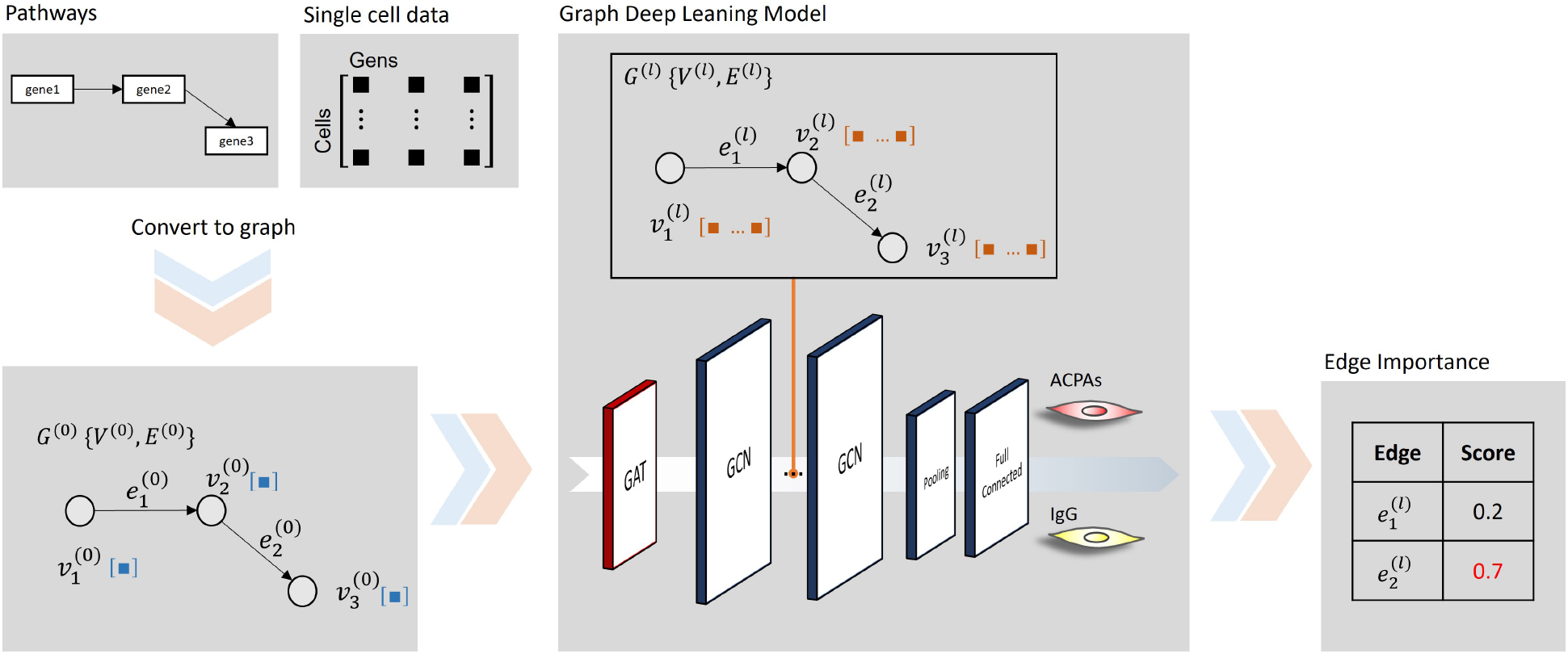
The framework to discovery signal pathway activation. The framework optimizes a GNN model to classify single cell data against ACPAs and IgG stimulated groups. Single cell data is profiled by RNA sequence from RA patients tissue derived FLS, which is represented to a graph data structure based on known pathways. Graph convolution layers are stacked to learn a deep representation of the input graph with respect to the classification. A graph attention layer is applied to learn the edges weights that indicate pathway activation.

In detail, each group has samples that are profiled by single cell sequencing. Each cell data can be presented as a graph that consists of nodes of genes and edges of signaling pathways. Denote a graph *G* = {*V, E*}, where *V* = {*υ_i_*}_*i*=1:*N^*υ*^*_ is the set of nodes, *N^υ^* is the total number of nodes and 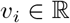 is the gene expression. *E* = {*e_i_*}_*i*=1:*N^e^*_ is the set of edges corresponding to the pathways. let *Y* = {*y_i_*}_*i*=1:*N^y^*_ labels the activation groups. The optimization aim in the framework is to learn a model Φ(*G*) that classifies *G* against *Y*.

The main component in the model is Graph Convolution Layer (GCN). Stacking that layers can learn a deep representation of the input graph with respect to the classification. Let *G^(0)^* be the input graph, and *G*^(*l*+1)^ be the representation after layer *l*. Equation 1 describes the computation applied in the layer *l*.

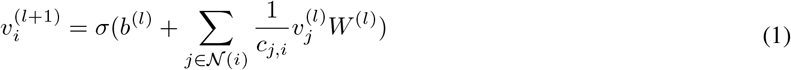

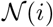 is the set of neighbors of node *i*. 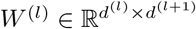 is the learnable weights of node attributes where *d^(l)^* are the user specified numbers. *C_j,i_* is the product of the square root of node degrees (i.e., 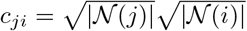), and *σ* is an activation function.

Graph Attention Layer (GAT) is a variant of GCN combined with the attention mechanism that checks the interactions between each node and its neighbors to evaluate which it should pay more attention to. This makes the model learn an attention score for each edge, which describes how they contribute differently to the results. The computation of GAT is described in Equation 2.

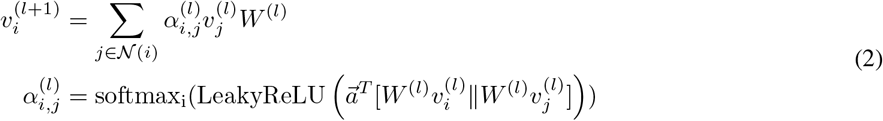

*a_i,j_* is the attention score associated with the edge between nodes *i* and *j*. It is computed by first concatenating the weighed *υ_i_*, *υ_j_*, 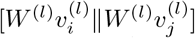, then takes a dot product of it and a learnable parameter vector 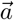 and activated by a LeakyReLU. Softmax is applied in the end to normalize the attention scores.

We apply one GAT directly on the input graph *G*^(0)^, followed by four GCN. *G^(k)^* output from the last GCN are aggregated in a pooling layer that squeeze *V^(k)^* to a vector 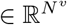, which is processed by a full-connected layer that feds the output to a Sigmoid function to produce a distribution over the 2 class labels.

## 4 Result

18 KEGG networks are selected as shown in Table 1, which includes 70 nodes and 139 edges. Their information is obtained from KEGG database and loaded by using Bio.KEGG.REST module of biopython Cock et al. [2009], which is then represented to graph data structure by using DGL, the Deep Graph Library Wang et al. [2019].

**Table 1:** Selected KEGG networks

| Network | Description |
| --- | --- |
| N00033 | EGF-EGFR-PI3K signaling pathway |
| N00030 | EGF-EGFR-RAS-PI3K signaling pathway |
| N00056 | Wnt signaling pathway |
| N00393 | ITGA/B-RhoGAP-RhoA signaling pathway |
| N01068 | ITGA/B-FAK-RAC signaling pathway |
| N01080 | ITGA/B-TALIN/VINCULIN signaling pathway |
| N00001 | EGF-EGFR-RAS-ERK signaling pathway |
| N00542 | EGF-EGFR-RAS-JNK signaling pathway |
| N00188 | IL1-IL1R-JNK signaling pathway |
| N00186 | IL1-IL1R-p38 signaling pathway |
| N01119 | RAC/CDC42-PAK-ERK signaling pathway |
| N00063 | TGF-beta signaling pathway |
| N00066 | MDM2-p21-Cell cycle G1/S |
| N00090 | p15-Cell cycle G1/S |
| N00091 | p27-Cell cycle G1/S |
| N00536 | MDM2-p21-Cell cycle G1/S |
| N00499 | ATR-p21-Cell cycle G2/M |
| N00493 | Spindle assembly checkpoint signaling |

The framework (Figure 1) is implemented in PyTorch Paszke et al. [2019] and DGL, which contains one GAT applied directly on the input graph; four GCN stacked sequentially; followed by an average pooling and a full-connected layer. The dimensional parameters *d^l^* of *W^(l)^* set to 1 and 50 for GAT and GCNs, respectively. ReLU is chosen as activation function *σ* for all GCN. And we add a Dropout layer with 0.1 drop rate after every GCN to prevent overfitting.

The dataset is divided to have 80% for training the model and rest 20% for the validation. The model is trained by using adaptive moment estimation (Adam) optimizer with a batch size of 200, and binary cross entropy is applied as the loss function. One training runs for 200 cycles through the training set, which takes 15 minutes on an NVIDIA Quadro RTX 4000 6GB GPU.

Training process was ran 10 times and their average performance is summarized. The classifier achieves 85% accuracy on both training and validation datasets. Based on the average attention score we arbitrarily selected edges that are weighted over 0.6 in Table 2 for further confirmation. In detail, FLS were stimulated with either ACPAs or IgG and cell lysate were collected for immunoblotting. Based on the edge importance, we investigate the top IL1-IL1R and bottom WNT-Frizzled signaling pathways, which are weighted higher than 0.6. In parallel, we also checked ITGA/B-PTK2 (FAK) pathway, which is weighted lower than 0.6.

**Table 2:**
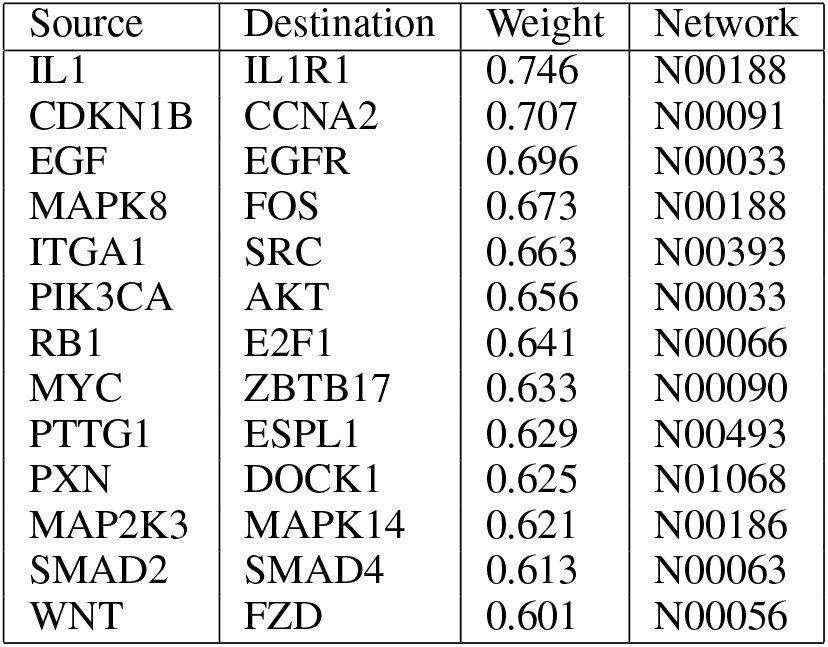
Edges Importance

| Source | Destination | Weight | Network |
| --- | --- | --- | --- |
| IL1 | IL1R1 | 0.746 | N00188 |
| CDKN1B | CCNA2 | 0.707 | N00091 |
| EGF | EGFR | 0.696 | N00033 |
| MAPK8 | FOS | 0.673 | N00188 |
| ITGA1 | SRC | 0.663 | N00393 |
| PIK3CA | AKT | 0.656 | N00033 |
| RB1 | E2F1 | 0.641 | N00066 |
| MYC | ZBTB17 | 0.633 | N00090 |
| PTTG1 | ESPL1 | 0.629 | N00493 |
| PXN | DOCK1 | 0.625 | N01068 |
| MAP2K3 | MAPK14 | 0.621 | N00186 |
| SMAD2 | SMAD4 | 0.613 | N00063 |
| WNT | FZD | 0.601 | N00056 |

In immunoblotting of kinetic study, protein expression of IL1R was up-regulated with presence of ACPAs in 2 hours and back to normal expression in 4 and 6 hours, whereas, regardless of time, IL1R expression remains no change with presence of IgG. (Figure 1A). The interleukin 1 receptor associated kinase 4 (IRAK4), which is in downstream event of IL1-IL1R signaling pathway, was also high expressed in 2 hours response to ACPAs stimulation (Figure 2A). In contrast, the expression of WNT5a/b, Frizzled5 and FAK were not affected with presence either ACPAs or IgG (Figure 2B).

**Figure 2:**
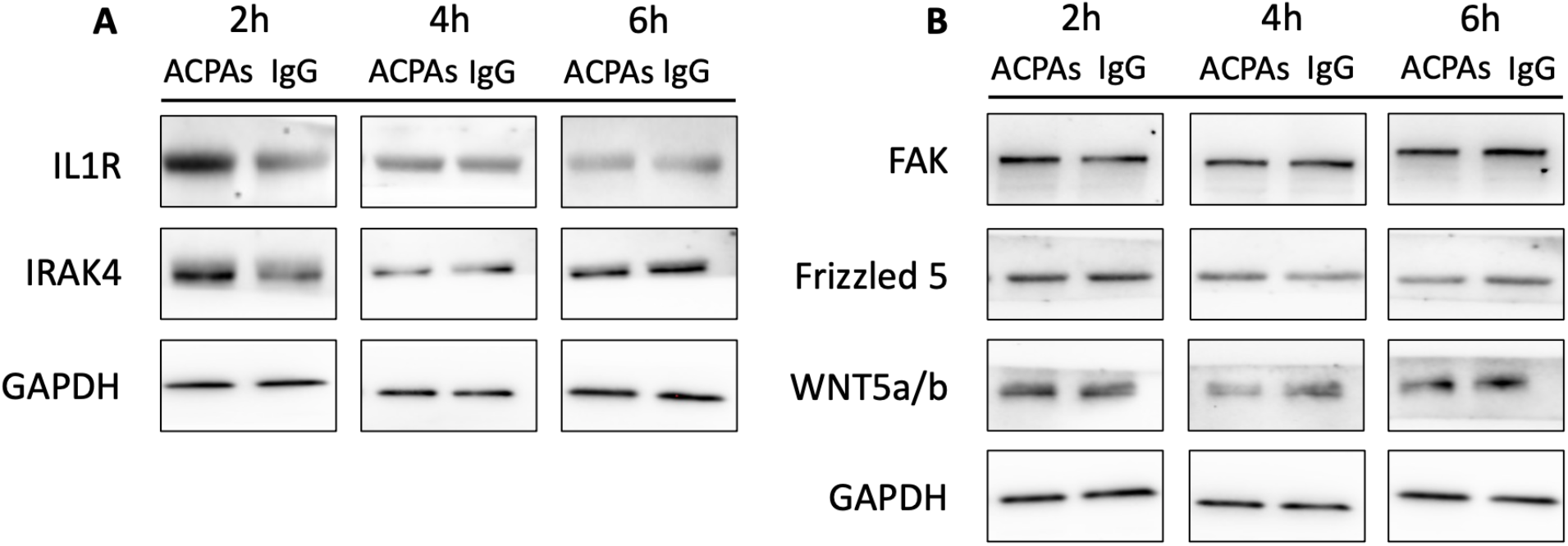
Activation of IL1-IL1R signaling pathway in ACPAs stimulated FLS. Starved FLS were exposed to 10 *μ*g/ml polyclonal ACPA IgG (ACPAs) or non-ACPA control IgG (IgG). Immunoblotting analysis of IL1R and IRAK4 are shown, whereas GAPDH protein was detected as control (A). The expression of protein FAK, Frizzled5 and WNT5a/b are analysed by immunoblotting, whereas GAPDH protein was detected as control (B). The presented data are representative for three independent experiments for all tested proteins.

## 5 Discussion

Investigation of signaling pathway in cells usually begin with dissecting observations, such as understanding morphologic changes, followed by locating potential signal tranduction candidates, which requests large amounts of literature search. Current experimental procedures take snapshots of signal transduction, such as phosphorylation, without the benefit of capture the whole stage and results are largely vary from expertness. In contrast, our study provides a method to investigate multiple signaling pathways simultaneously. The graph deep learning model searched in 18 networks with 139 signal transduction candidates, from which the found pathway weights effectively facilitate experimental confirmation.

The highlights of this study is both new biologic findings, and the successful application of the new graph deep learning approach to assessing single-cell signaling from sc-mRNA sequencing data. This approach holds promise beyond RA and ACPA-stimulated cells and can be generalized ot other cell signaling pathways, using a similar data driven approach and experimental confirmation. We demonstrated that the top weighted genes interleukin 1 receptor (IL1R) was up-regulated in protein expression in ACPA-stimulated FLS. IL1R is a cytokine receptor that binds interleukin 1 (IL1), which is one of the most important pro-inflammatory cytokines in the pathophysiology of RA. The activation of IL1-IL1R signaling has immune and pro-inflammatory action and correlates with disease activity, progression of joint destruction of RA Dinarello [2018]. Despite it has been reported that ACPA are able to bind PBMC and up-regulate IL-1beta mRNA expression level compared with control IgG Gertel et al. [2018], our current report is the first to investigate the potential relationship between IL1R and ACPAs in RA FLS.

Meanwhile, IL-1 binds first to IL-1R1, that recruits the IL-1 receptor accessory protein (IL-1RAcP), which serves as a co-receptor. After the formation of receptor heterodimeric complex, two intracellular adaptor proteins are assembled, one of which is interleukin-1 receptor-activated protein kinase 4 (IRAK4). To look insight, we also evaluated protein expression of IRAK4. Immunoblotting of IRAK4 illustrated an elevated protein expression in ACPA-stimulated FLS. Taken together, these results suggest a novel finding that ACPAs up-regulate IL1-IL1R pathway that potential enhance inflammation.

In addition, the activation of PI3K signaling pathway is also ranked with high weight, which is in line with our previous finding Sun et al. [2019] that shows a role for PI3K/Akt in regulating ACPA-induced migration of challenged FLS. PI3K modulates organization of actin cytoskeleton and lamellipodium formation via Akt phosphorylation, which can result in cytoskeleton remodeling and increased cell motility.

We also selected several low weighed nodes for further confirmation. As we expected, the expression of FAK, Fizzled5 and WNT5a/b remains no change in all time points of both groups. These results suggest that FAK adhesion and Wnt signaling pathway are inactivated regardless of stimulation.

Unlike commonly used stimulus, ACPAs consist of a group of antibodies with different specificities that target various citrullinated proteins/peptides Demoruelle and Deane [2011]. ACPAs have been suggested to contribute to joint pain and bone loss already before onset of joint inflammation in RA Bos et al. [2010], Kleyer et al. [2014], Krishnamurthy et al. [2019], Rantapää-Dahlqvist et al. [2003]. Further, ACPAs with different specificities binds to surface of developing osteoclasts (OC) and increase OC differentiation and bone erosion in vitro and induce pain behaviors in mouse model Harre et al. [2012], Krishnamurthy et al. [2016], Wigerblad et al. [2016]. Together with our report, all evidence suggest that ACPAs contribute to multiple symptoms, through different mechanisms.

As every other method, it should be mentioned that there are certain limitations in this study needed to be investigated in future. The dataset used in this study is small, while the deep learning model usually need large volume of data for training. We didn’t distinguish edge types in the model, however signaling pathway consists of different interaction types, such as expression, inhibition, etc. Validating the proposed framework on larger datasets with improved model and data structure will be planed. While we selected top ranked edges for experimental confirmation based on an arbitrary threshold, a precise method to define the threshold and interpret the ranks remains to future works.

In conclusion, we proposed a graph deep learning framework that efficiently identifies activated signaling pathways from sc-mRNA-seq data. Based on this, we describe a novel potential pathogenic mechanisms of ACPAs in RA. Considering the diverse clinic presentations among ACPA positive RA patients and a body of accumulated evidence in research, the broad specificities in polyclonal ACPA contribute to distinct pathogenic events, potentially through activation of multiple parallell signalling pathways. Therefore, the mechanism of action in ACPA triggered signaling transduction is important for better understanding the pathogenisis of RA, and its diversity among patients.

## Acknowledgements

M.S, Y.C and P.B designed and conceptualized the method. M.S performed the experiments and analyse. Y.C wrote all the code and implemented the methods. M.S and Y.C wrote manuscript with input of other authors. M.S and Y.C contributed equally to this work.

